# Early life gut microbial and metabolic shifts reflected in stool consistency- a gut transit time proxy

**DOI:** 10.1101/2024.06.05.597641

**Authors:** Anna-Katariina Aatsinki, Matilda Kråkström, Katja Salonen, Heidi Isokääntä, Abhijit Paul, Thomaz Bastiaanssen, Leo Lahti, Minna Lukkarinen, Eveliina Munukka, Tiina Paunio, Katri Kantojärvi, Pieter C. Dorrestein, Hasse Karlsson, Linnea Karlsson, Matej Oresic, Alex M Dickens, Santosh Lamichhane

**Affiliations:** Centre for Population Health Research, University of Turku and Turku University Hospital, Turku, Finland; FinnBrain Birth Cohort Study, Turku Brain and Mind Center, Department of Clinical Medicine, University of Turku, Turku, Finland; Turku Bioscience Centre, University of Turku and Åbo Akademi University, 20520 Turku, Finland; Research Center for Infections and Immunity, Institute of Biomedicine, University of Turku,Turku, Finland; Amsterdam University Medical Center, Amsterdam, Netherlands; Department of Computing, University of Turku, Turku, Finland; Department of Pediatrics and Adolescent Medicine, Turku University Hospital and University of Turku, Turku, Finland; Turku Clinical Microbiome Bank, Clinical Microbiology, Turku University Hospital, The Wellbeing services county of Southwest Finland, Turku, Finland; Department of Psychiatry and SleepWell Research Program, Faculty of Medicine, University of Helsinki, Helsinki University Central Hospital, Helsinki, Finland; Department of Public Health, University of Turku, Turku, Finland; Skaggs School of Pharmacy and Pharmaceutical Sciences, University of California San Diego, La Jolla, CA, USA; Department of Psychiatry, University of Turku and Turku University Hospital, Turku, Finland; Department of Public Health, University of Turku, Turku, Finland; Department of Child Psychiatry, Turku University Hospital Turku, Finland; School of Medical Sciences, Örebro University, 702 81 Örebro, Sweden; Department of Chemistry, University of Turku, 20520 Turku, Finland; Microbe Centre, University of Turku and Wellbeing Services County of Southwest Finland, Turku, Finland

## Abstract

Transit time, fluid intake, diet, and overall gut health influence stool consistency. However, the relationships between the gut metabolome (microbial metabolites) and stool consistency in infants and young children remain poorly understood. Here, we analyzed the metabolome and microbiota of 618 stool samples from children aged 2.5 months (n = 360), 6 months (n = 229), 14 months (n = 274), and 30 months (n = 169) from the FinnBrain Birth Cohort Study, and related these data to stool water content and parent-reported stool consistency. Breastfeeding showed the strongest association with both stool consistency and fecal water content. Concentrations of newly identified microbial bile acid amidates were associated with constipation and stool water content, while bile salt hydrolase, an enzyme involved in bile acid deconjugation and conjugation was predicted to be negatively associated with stool water content. In addition, short-chain fatty acids, particularly acetate, were positively associated with stool water content, whereas branched-chain short-chain fatty acids showed negative associations. These findings suggest that longer gut transit time permits more extensive microbial metabolism, including the transformation of bile acid amidates. We also found that microbial taxonomic richness, diversity, and community composition were primarily associated with stool water content, with only weak associations with stool consistency. Overall, our results highlight the importance of documenting stool consistency in fecal metabolomics and microbiome research, and provide new insights into how breastfeeding, microbial metabolism, and gut transit time shape early-life gut development.

## Introduction

The concept of assessing stool consistency and its link with gastrointestinal pathophysiology dates back over a century [1]. These days, stool consistency is generally assessed using the Bristol Stool Scale. Recent reports suggest that stool consistency is strongly associated with gut microbiota richness, composition, and microbial metabolism [2–5] . Additionally, pathological conditions related to alterations in stool consistency or defecation frequency, such as chronic constipation, are associated with gut microbiota composition in children [6].

Emerging literature, including our own work, suggests that the gut microbiota in early life plays a pivotal role in host health and the development of various diseases across the lifespan [7, 8] . There is a growing awareness that the impact of microbes on various physiological processes including initiation and pathogenesis of various diseases may be mediated by microbially regulated metabolites [9–12]. Current microbiota research related to human health and disease has shifted its focus from characterizing community composition towards understanding the intricate interplay between the gut microbiota and microbiota-dependent metabolites. Efforts have been made to enhance the identification of microbiota composition [2, 13–16]. Accounting for these factors will enable us to more robustly identify disease markers in human microbiota studies [17]. Particularly high intra-individual variance in gut microbiota community composition is explained by proxies for gut transit time [18] . However, measuring the gut transit time is often inconvenient due to logistic challenges and its invasive nature [16, 19]. Stool consistency measurement has been identified as a potentially useful proxy for gut transit time within specific cohorts [20]; although other explanations are possible - in general hard stools suggest slow transit while loose stools are linked with fast transit through the colon.

To the best of our knowledge, there are no studies in children investigating stool metabolites in relation to stool consistency. Several adult human studies from both healthy and pathological conditions have identified associations with gut microbiota composition, stool consistency, or transit time measurement [17]. Although understanding in adult populations is increasing, there is a dearth of studies examining the impact of the gut microbiota and metabolome on both stool consistency and total water content of feces or *vice versa* in children *[21]*. Here, we aim to characterize associations between stool consistency, total water content of stool, stool microbiota composition, and metabolite levels in infants and small children. We used stool samples and complementary markers for gut transit time: Stool water content and parental reports of crude stool consistency from 2.5, 6, 14, and 30-month-old children.

## Material and methods

### Cohort and Questionnaire Data

The study subjects are children from the FinnBrain Cohort Study [22] that is a general population birth cohort study located in the southwestern Finland. The Ethical Committee of Southwestern Finland approved the study. Parents provided informed consent on behalf of their children.

The sample used in the current study were a subset of children that participated in a study visit at 2.5, 6, 14 and 30 months postpartum. The fecal samples were collected from May 2013 to May 2018. The samples were collected in plastic tubes, and parents were instructed to store the sample in a refrigerator, and bring the sample to the laboratory within 24 h. The sample collection time was reported.

Information on breastfeeding was collected with questionnaires to the mother at 3, 6, 12, and 24 months postpartum and during study visits 2.5, 6, 14, and 30 months. Information on delivery mode and gestational age was collected from the hospital records (The Wellbeing Services County of Southwest Finland, VARHA). Breastfeeding was categorized as any current breastfeeding (yes vs. no). At the sample collection, the parent reported the child stool consistency as: normal, loose, constipated, gas or other.

### DNA Extraction, Sequencing and Data Processing

The samples were divided into cryotubes and freezed in -80C within 2 days after arriving at the laboratory. Samples were kept at +4C before freezing. Only samples that were freezed within 48 h of sample collection were sequenced. DNA was extracted using a semi-automatic extraction instrument Genoxtract with DNA stool kit (HAIN life science, Germany). The DNA extraction was performed according to the manufacturer’s protocol except that 1 ml of lysis buffer was added and the samples were homogenized with glass beads 1000 rpm / 3 min. The samples were centrifuged at high speed (> 13000 rpm) for 5 min. The lysate (800μL) was then transferred to tubes and the extraction proceeded according to the manufacturer’s protocol. DNA yields were measured with Qubit fluorometer using Qubit dsDNA High Sensitivity Assay kit (Thermo Fisher Scientific, USA). Bacterial community composition was determined by sequencing the V4 region of 16S rRNA gene using Illumina MiSeq platform (Illumina, USA). The sequence library was constructed with an in-house developed protocol as described in [23, 24].Positive control (DNA 7-mock standard) and negative control (PCR grade water) were included in library preparation and sequencing runs.

DADA2-pipeline (version 1.14) was used to preprocess the 16S rRNA gene sequencing data to infer exact amplicon sequence variants (ASVs)[25]. The reads were truncated to length 225 and reads with more than two expected errors were discarded (maxEE = 2). SILVA taxonomy database (version 138) and RDP Naive Bayesian Classifier algorithm ref were used for the taxonomic assignments of the ASVs [26].

### Stool Water Content

To quantify the water content, stool samples underwent freeze-drying. Prior drying, samples were weighted and briefly centrifuged in the bottom of the tubes. Subsequently, they were transferred to a custom freeze-drier equipped with a vacuum pump. The samples were left to dry overnight and then reweighed to determine their dry weight.

### Metabolite Analyses

The BAs, SCFA and untargeted metabolome were previously described and adapted for this publication[23, 27].

### Targeted BAs measurement

Conjugated BA amidates were extracted by adding 80 μL fecal homogenate to 400 μL crash solvent (methanol containing 62,5 ppb each of the internal standards TCA-d4, GUDCA-d4, GCA-d4, CA-d4, UDCA-d4, GCDCA-d4, CDCA-d4, DCA-d4, GLCA-d4 and 625 ppb of LCA-d4) and filtering them using a Supelco protein precipitation filter plate after vortexing. The samples were dried under a gentle nitrogen flow and resuspended using 40 μL resuspension solution (Methanol:water (40:60) as in injection standard). Quality control (QC) samples were prepared by combining an aliquot of every sample into a tube, vortexing it and preparing QC samples in the same way as the other samples. Blank samples and conjugated BA standard mixture (made initially by combinatorial chemistry) were prepared by pipetting 400 μL crash solvent into a 96-well plate, then drying and resuspending them the same way as the other samples.

The LC separation was performed on an Acquity Premier system consisting of a binary pump, an autosampler set to 10 °C and a column oven set to 45 °C. An Acquity premier HSS T3 1.8um (2.1x100 mm) column with a precolumn with the same material was used. Eluent A was 2 mM ammonium acetate in water and eluent B was 2 mM ammonium acetate in methanol.

The gradient started with 33.5 % B for half a minute and increased to 72 % B over three minutes. Next, the gradient increased to 100 % B over 7.5 minutes and was kept at 100 % B for 5 minutes. The flow rate was 0.4 mL/min and the injection volume was 5 μL. The mass spectrometer used for this method was a Sciex 7600 ZenoTof mass spectrometer operating in information-dependent mode with positive polarity. The ion source gas 1 was 60 psi and gas 2 was 80 psi. The curtain gas was 35 psi, the CAD gas was 7 and the temperature was 450°C. The spray voltage was 5500 V.

The BAs were measured in fecal samples as described previously [23, 28]. Only samples frozen within 24 h of sample collection were included in the metabolome analyses. The order of the samples was randomized before sample preparation. Two aliquots (50 mg) of each fecal sample were weighed. An aliquot was freeze-dried prior to extraction to determine the dry weight. The second aliquot was homogenized by adding homogenizer beads and 20 μL of water for each mg of dry weight in the fecal sample, followed by samples freezing to at least -70 °C and homogenizing them for five minutes using a bead beater. The BAs analysed were Litocholic acid (LCA), 12-oxo-litocholic acid(12-oxo-LCA), Chenodeoxycholic acid (CDCA), Deoxycholic acid (DCA), Hyodeoxycholic acid (HDCA), Ursodeoxycholic acid (UDCA), Dihydroxycholestanoic acid (DHCA), 7-oxo-deoxycholic acid (7-oxo-DCA), 7-oxo-hyocholic acid (7-oxo-HCA), Hyocholic acid(HCA), β-Muricholic acid (b-MCA), Cholic acid (CA), Ω/α-Muricholic acid (w/a-MCA), Glycolitocholic acid (GLCA), Glycochenodeoxycholic acid (GCDCA), Glycodeoxycholic acid (GDCA), Glycohyodeoxycholic acid (GHDCA), Glycoursodeoxycholic acid (GUDCA), Glycodehydrocholic acid (GDHCA), Glycocholic acid (GCA), Glycohyocholic acid (GHCA), Taurolitocholic acid (TLCA), Taurochenodeoxycholic acid (TCDCA), Taurodeoxycholic acid (TDCA), Taurohyodeoxycholic acid (THDCA), Tauroursodeoxycholic acid (TUDCA), Taurodehydrocholic acid (TDHCA), Tauro-α-muricholic acid (TaMCA), Tauro-β-muricholic acid (TbMCA), Taurocholic acid (TCA), Trihydroxycholestanoic acid (THCA) and Tauro-Ω-muricholic acid (TwMCA). BAs were extracted by adding 40 μL fecal homogenate to 400 μL crash solvent (methanol containing 62,5 ppb each of the internal standards LCA-d4, TCA-d4, GUDCA-d4, GCA-d4, CA-d4, UDCA-d4, GCDCA-d4, CDCA-d4, DCA-d4 and GLCA-d4) and filtering them using a Supelco protein precipitation filter plate. The samples were dried under a gentle flow of nitrogen and resuspended using 20 μL resuspenstion solution (Methanol:water (40:60) with 5 ppb Perfluoro-n-[13C9]nonanoic acid as in injection standard). Quality control (QC) samples were prepared by combining an aliquot of every sample into a tube, vortexing it and preparing QC samples in the same way as the other samples. Blank samples were prepared by pipetting 400 μL crash solvent into a 96-well plate, then drying and resuspending them the same way as the other samples. Calibration curves were prepared by pipetting 40 μL of standard dilution into vials, adding 400 μL crash solution and drying and resuspending them in the same way as the other samples. The concentrations of the standard dilutions were between 0.0025 and 600 ppb. The LC separation was performed on a Sciex Exion AD 30 (AB Sciex Inc., Framingham, MA) LC system consisting of a binary pump, an autosampler set to 15 °C and a column oven set to 35 °C. A waters Aquity UPLC HSS T3 (1.8μm, 2.1×100mm) column with a precolumn with the same material was used. Eluent A was 0.1 % formic acid in water and eluent B was 0.1 % formic acid in methanol. The gradient started from 15 % B and increased to 30 % B over 1 minute. The gradient further increased to 70 % B over 15 minutes. The gradient was further increased to 100 % over 2 minutes. The gradient was held at 100 % B for 4 minutes then decreased to 15 % B over 0.1 minutes and re-equilibrated for 7.5 minutes. The flow rate was 0.5 mL/min and the injection volume was 5 μL. The mass spectrometer used for this method was a Sciex 5500 QTrap mass spectrometer operating in scheduled multiple reaction monitoring mode in negative mode. The ion source gas1 and 2 were both 40 psi. The curtain gas was 25 psi, the CAD gas was 12 and the temperature was 650 °C. The spray voltage was 4500 V. Data processing was performed on Sciex MultiQuant.

### Quantification of SCFA

We adapted and modified the targeted SCFA analysis from previous work [29, 30]. Fecal samples were homogenized by adding water (10 μL per mg of dry weight as determined for the BA analysis) to wet feces, the samples were homogenized using a bead beater. Analysis of SCFA was performed on fecal homogenate (50 μL) crashed with 500 μL methanol containing internal standard (propionic acid-d6 and hexanoic acid-d3 at 10 ppm). Samples were vortexed for 1 min, followed by filtration using 96-Well protein precipitation filter plate (Sigma-Aldrich, 55263-U). Retention index (RI, 8 ppm C10-C30 alkanes and 4 ppm 4,4-Dibromooctafluorobiphenyl in hexane) was added to the samples. Gas chromatography (GC) separation was performed on an agilent 5890B GC system equipped with a Phenomenex Zebron ZB-WAXplus (30 m × 250 μm × 0.25 μm) column a short blank pre-column (2 m) of the same dimensions was also added. A sample volume of 1 μL was injected into a split/splitless inlet at 285°C using split mode at 2:1 split ratio using a PAL LSI 85 sampler. Septum purge flow and split flow were set to 13 mL/min and 3.2 mL/min, respectively. Helium was used as carrier gas, at a constant flow rate of 1.6 mL/min. The GC oven program was as follows: initial temperature 50°C, equilibration time 1 min, heat up to 150°C at the rate of 10°C/min, then heat at the rate of 40°C/min until 230°C and hold for 2 min. Mass spectrometry was performed on an Agilent 5977A MSD. Mass spectra were recorded in Selected Ion Monitoring (SIM) mode. The detector was switched off during the 1 min of solvent delay time. The transfer line, ion source and quadrupole temperatures were set to 230, 230 and 150°C, respectively. Dilution series of SCFA standards of acetic, propionic, butyric, valeric, hexanoic acid, isobutyric, and iso-valeric acid were prepared in concentrations of 0.1, 0.5, 1, 2, 5, 10, 20, 40, and 100 ppm for the construction of standard curves for quantification. SCFAs were quantified using Agilent Masshunter.

### Polar metabolites were extracted in methanol

The method was adapted from the method used by [31]. Fecal homogenate (60 μL) was diluted with 600 μL methanol crash solvent containing internal standards (heptadecanoic acid (5 ppm) valine-d8 (1 ppm) and glutamic acid-d5 (1 ppm)). After precipitation the samples were filtered using Supelco protein precipitation filter plates. One aliquot (50 μL) was transferred to a shallow 96-well plate to create a QC sample. The rest of the sample volume was dried under a gentle stream of nitrogen and stored in -80 °C until analysis. After thawing the samples were again dried to remove any traces of water. Derivatization was carried out on a Gerstel MPS MultiPurpoe Sampler using the following protocol: 25 μL methoxamine (20 mg/mL) was added to the sample followed by incubation on a shaker heated to 45 °C for 60 minutes. N-Methyl-N-(trimethylsilyl) trifluoroacetamide (25 μL) was added followed by incubation (60 min). After that, 25 μL retention index was added, the sample was allowed to mix for one min followed by injection. The automatic derivatization was carried out using the Gerstel maestro 1 software (version 1.4). Gas chromatographic (GC) separation was carried out on an Agilent 7890B GC system equipped with an Agilent DB-5MS (20 m x 0,18 mm (0,18 μm)) column. A sample volume of 1 μl was injected into a split/splitless inlet at 250°C using splitless mode. The system was guarded by a retention gap column of deactivated silica (internal dimensions 1.7 m, 0.18 mm, PreColumn FS, Ultimate Plus Deact; Agilent Technologies, CA, USA). Helium was used as carrier gas at a flow rate of 1.2 ml/min for 16 min followed by 2 mL/min for 5.75 min. The temperature programme started at 50°C (5 min), then a gradient of 20°C/min up to 270°C was applied and then finally a gradient of 40°/min to 300°C, where it was held stable for 7 min. The mass spectrometry was carried out on a LECO Pegasus BT system (LECO). The acquisition delay was 420 sec. The acquisition rate was 16 spectra/sec. The mass range was 50 – 500 m/z and the extraction frequency was 30 kHz. The ion source was held at 250 °C and the transferline heater temperature was 230 °C. ChromaTOF software (version 5.51) was used for data acquisition. The samples were run in 9 batches, each consisting of 100 samples and a calibration curve. In order to monitor the run a blank, a QC and a standard sample with a known concentration run between every 10 samples. Between every batch the septum and liner on the GC were replaced, the precolumn was cut if necessary and the instrument was tuned. The retention index was determined with ChromaTOF using the reference method function. For every batch a reference file was created. The reference file contained the spectras and approximate retention times of the alkanes from C10 to C30 as determined manually). A reference method was implemented for every sample in order to determine the exact retention time of the alkanes. Text files with the names and retention times of the alkanes were then exported and converted to the correct format for MSDIAL using an in-house R script. The samples were exported from ChromaTOF using the netCDF format. After this they were converted to abf files using the abfConverter software (Reifycs). Untargeted data processing was carried out using MSDIAL (version 4.7). The minimum peak height was set to an amplitude of 1000, the sigma window value was 0.7 and the EI spectra cut off was 10. The identification was carried out using retention index with the help of the GCMS DB-Public-kovatsRI-VS3 library provided on the MSDIAL webpage. A separate RI file was used for each sample. The RI tolerance was 20 and the m/z tolerance was 0.5 Da. the EI similarly cut off was 70 %. The identification score cut off was 70 % and retention information was used for scoring. Alignment was carried out using the RI with an RI tolerance of 10. The EI similarity tolerance was 60 %. The RI factor was 0.7 and the EI similarity factor was 0.5. The results were exported as peak areas and further processed with excel. In excel the results were normalized using heptadecanoic acid as internal standard and the features with a coefficient of variance of less than 30 % in QC samples were selected. Further filtering was carried out to remove alkanes and duplicate features. The IDs of the features which passed the CV check were further checked using the Golm Metabolome Database.

### Predicted functional potential of microbiome

We used PICRUSt2 for predicting community functions[32]. We used reference sequences and count abundances of detected ASVs with abundance more than 0.1/100 and prevalence more than 2/100. We used default settings in PICRUSt2 version 2.5.0. Then, for downstream analyses, we focused on enzymes predicted by Enzyme Commission (EC) numbers involved in metabolizing short-chain fatty acids or bile acids, as these metabolites are robustly associated with water content or stool consistency. Relevant ECs were identified based on substrate and product information available on KEGG [33].

### Polygenic score for defecation frequency

DNA was extracted from cord blood samples and DNA was sequenced at the National Institute for Health and Welfare and genotyped with Illumina Infinium PsychArray and Illumina Infinium Global Screening Array at the Estonian Genome Centre as previously described [34]). Genotyped data was pre-phased with Eagle 2.3.5 [35]and imputed with Beagle 4.1 [36]using the Finnish population-specific SISu v2 imputation reference panel. Polygenic score for defecation frequency was calculated based on[37]. PGS was computed using the PRS-CS method [38]with Bayesian regression to estimate posterior variant effect sizes, employing the European 1000 Genomes Project samples as a reference panel. The final PGS for each participant was calculated as a weighted sum of about one million risk allele counts and standardized to a mean of 0 and a standard deviation of 1 using R 3.6.0.

### Statistical analyses

All of the analyses were conducted, and figures created in R (version 4.3.2) using packages mia (version 1.10.2), miaViz (version 1.10.2), scater (version 1.30.1), vegan (version 2.6.2004), nlme (version 3.1.164) and tidyverse (version 2.0.0) [39–42]. The markers of stool consistency were modeled with: 1. water content as continuous variable, 2. stool consistency as a 4-class variable, 3. PGS as continuous variable. All analyses were conducted with all the timepoints as well as separately for each age group.

To describe the longitudinal pattern of reported stool consistency, binary variables of a. loose vs no loose, b. constipation vs no constipation, c. gas and other complaints vs no were created. Subsequently, differences between timepoints were tested with Kruskal-Wallis and Dunn’s posthoc test. To inform the interconnectedness of the stool consistency and water content as well as background factors, Spearman correlation, Wilcoxon signed rank test as well as Kruskal-Wallis test were used.

Metabolome data was log2-transformed with half of the smallest value as a pseudocount. Associations with water content were tested with Spearman correlation. Associations with stool consistency were tested with Kruskal-Wallis test. The longitudinal data was tested with linear mixed model with metabolite concentration as the dependent variable, stool consistency or water content as fixed effects, and subject as the random effect. PGS was modelled similarly except without breastfeeding as a covariate. The age terms were fitted as splines with break points at 6 and 14 months. These correspond to the age of solid food introduction and the age when all food groups should be present in the diet according to the local nutrition recommendations.

All EC abundances from PICRUSt2 were transformed to relative abundances and further log-transformed. The subset of ECs involved in SCFA or BA metabolism were selected for downstream testing. The associations were calculated with Spearman correlation for water content and Kruskal-Wallis test for the stool consistency variables. Dunn’s post-hoc test was calculated following significant Kruskal-Wallis test. Linear regression models with EC abundance as dependent variable, water content and breastfeeding as independent variables were built per time point. The longitudinal analyses were conducted with mixed models with EC abundance as the dependent variable, breastfeeding and water content or stool consistency as the independent variables and subject identity as the random effect. P-values were corrected for multiple testing with Benjamini-Hochberg, and we consider q<0.05 as statistically significant.

Taxonomic diversity, i.e. alpha diversity, was estimated with Shannon diversity and observed richness (number of observed species) from non-transformed ASV-level data with the function addAlpha from the mia package, and 10 iterations were used for the rarefication.

The longitudinal analyses were conducted with mixed models with alpha diversity as the dependent variable, breastfeeding and water content, PGS or stool consistency as the independent variables and subject as the random effect. Community composition i.e. beta diversity, was tested with distance-based redundancy analysis (db-RDA) with non-rarefied relative abundance ASV-level data and Bray-Curtis dissimilarity and Aitchinson distances.

Differentially abundant genera (>0.1 prevalence) were tested with MaAsLin 3 with the following model: abundance ∼ stool water content / stool consistency + current breastfeeding + age + (1 | subject). PGS was also modelled similarly with the expectation that breastfeeding was not included as a covariate. Total sum scaling normalization and log-transformation were applied.

## Results

### Longitudinal measurements

We measured the gut microbiota, metabolome, stool consistency, and water content from 618 stool samples (Figure 1A). We obtained measurements at four time points of 2.5 months (n=360), 6 months (n=229), 14 months (n=274), and 30 months (n=169) of age. The distribution of participants providing samples was as follows: those who provided only one sample (n=326), two samples (n=190), three samples (n=82), and all four samples (n=20). We performed targeted metabolomics analysis of six short-chain fatty acids (SCFAs; acetic, propionic, iso-butyric, butyric, isovaleric, and valeric acids), and bile acids (BAs; including both taurine and glycine conjugates (n=33) and those microbial conjugated to 22 different amino acids (n=110)). We also detected metabolites panel including a wide range of chemical classes, such as amino acids, carboxylic acids (mainly free fatty acids and other organic acids), hydroxy acids, phenolic compounds, alcohols, and sugar derivatives in an untargeted metabolomics assay. Parents reported on crude stool consistency. Stool water content was measured as the percentage of water present in the stool sample. Table 1 shows clinical characteristics of the study subjects and Table S1 shows the sample sizes for available data types including measured metabolites, stool consistency, and water content analyses.

**Figure 1.**
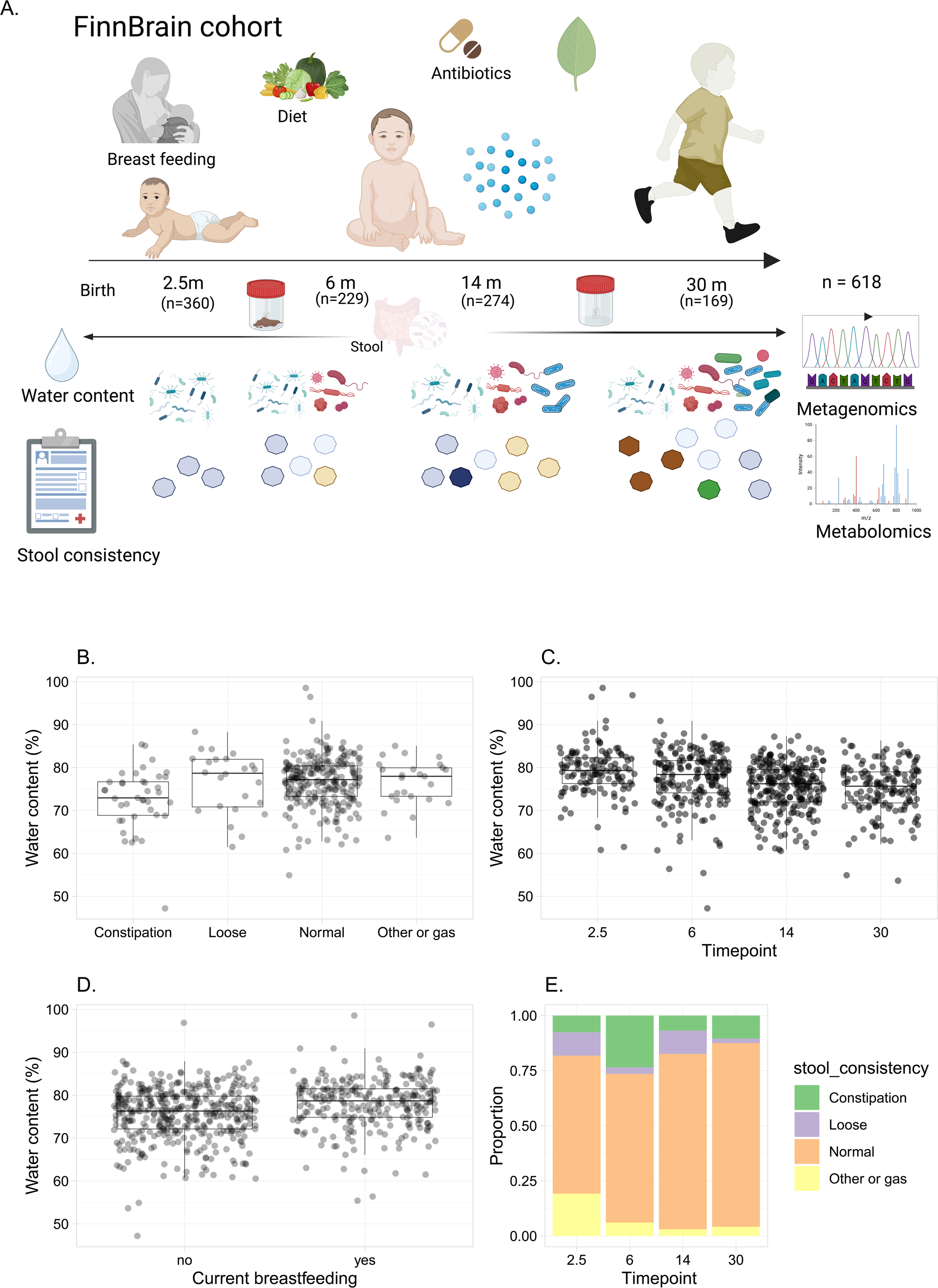
A. An overview of the study setting. We obtained longitudinal gut microbiota (16S rRNA), metabolome (targeted (bile acid, SCFAs) and untargeted (polar metabolites)), stool consistency, and water content measurements at four time points of 2.5 months (n=360), 6 months (n=229), 14 months (n=274), and 30 months (n=169) of age. Short-chain fatty acids (SCFAs). The figure is made in Biorender. B. Constipated subjects had lower water content. Water content gradually decreased as the child aged, and consequently water content was associated with breastfeeding. D. There were more loose stools and reports of gas and other complaints at 2.5 months, whereas constipation was reported more at 6 months.

**Table 1.** Characteristics of the study subjects who had either stool consistency or water content were reported.

| Timepoint <u>(months)</u> | 2.5 | 6 | 14 | 30 | Overall |
| --- | --- | --- | --- | --- | --- |
| Sample sizes | 360 | 229 | 274 | 169 | 1032 |
| Stool consistency <u>(number of individuals, % of the total sample)</u> |  |  |  |  |  |
| constipation | 23 (6.4%) | 31 (13.5%) | 9 (3.3%) | 10 (5.9%) | 73 (7.1%) |
| loose | 32 (8.9%) | 4 (1.7%) | 14 (5.1%) | 2 (1.2%) | 52 (5.0%) |
| normal | 189 (52.5%) | 89 (38.9%) | 105 (38.3%) | 80 (47.3%) | 463 (44.9%) |
| other or gas | 58 (16.1%) | 8 (3.5%) | 4 (1.5%) | 4 (2.4%) | 74 (7.2%) |
| missing | 58 (16.1%) | 97 (42.4%) | 142 (51.8%) | 73 (43.2%) | 370 (35.9%) |
| Water content <u>(%)</u> |  |  |  |  |  |
| mean (SD) | 79.3 (5.70) | 77.4 (6.43) | 75.4 (5.50) | 75.0 (5.86) | 76.6 (6.06) |
| median [Min, Max] | 79.3 [60.8, 98.6] | 78.4 [47.2, 90.9] | 76.2 [60.5, 87.3] | 75.7 [53.6, 86.3] | 77.2 [47.2, 98.6] |
| missing | 236 (65.6%) | 42 (18.3%) | 30 (10.9%) | 23 (13.6%) | 331 (32.1%) |
| Delivery mode |  |  |  |  |  |
| C-section | 57 (15.8%) | 37 (16.2%) | 45 (16.4%) | 23 (13.6%) | 162 (15.7%) |
| vaginal | 298 (82.8%) | 191 (83.4%) | 226 (82.5%) | 145 (85.8%) | 860 (83.3%) |
| missing | 5 (1.4%) | 1 (0.4%) | 3 (1.1%) | 1 (0.6%) | 10 (1.0%) |
| Gestational age <u>(weeks)</u> |  |  |  |  |  |
| mean (SD) | 40.1 (1.36) | 40.0 (1.38) | 39.8 (1.52) | 39.9 (1.36) | 40.0 (1.41) |
| median [Min, Max] | 40.1 [34.6, 42.3] | 40.0 [34.1, 42.3] | 40.1 [32.9, 42.3] | 40.0 [35.6, 42.3] | 40.1 [32.9, 42.3] |
| Antibiotic treatment prior 6 months |  |  |  |  |  |
| no | 300 (83.3%) | 128 (55.9%) | 111 (40.5%) | 75 (44.4%) | 614 (59.5%) |
| yes | 2 (0.6%) | 4 (1.7%) | 22 (8.0%) | 21 (12.4%) | 49 (4.7%) |
| missing | 58 (16.1%) | 97 (42.4%) | 141 (51.5%) | 73 (43.2%) | 369 (35.8%) |
| Current <u>breastfeeding</u> |  |  |  |  |  |
| full breastfeeding | 220 (61.1%) | 9 (3.9%) | 0 (0%) | 0 (0%) | 229 (22.2%) |
| no breastfeeding | 7 (1.9%) | 1 (0.4%) | 3 (1.1%) | 1 (0.6%) | 12 (1.2%) |
| partial<br>breastfeeding | 50 (13.9%) | 145 (63.3%) | 32 (11.7%) | 1 (0.6%) | 228 (22.1%) |
| ceased<br>breastfeeding | 10 (2.8%) | 53 (23.1%) | 217 (79.2%) | 155 (91.7%) | 435 (42.2%) |
| missing | 73 (20.3%) | 21 (9.2%) | 22 (8.0%) | 12 (7.1%) | 128 (12.4%) |
| Maternal age (years) |  |  |  |  |  |
| mean (SD) | 30.8 (4.42) | 30.7 (4.52) | 31.1 (4.33) | 31.0 (4.29) | 30.9 (4.39) |
| median [Min, Max] | 31.0 [19.0,<br>45.0] | 30.0 [19.0,<br>42.0] | 31.0 [19.0,<br>45.0] | 30.0 [21.0,<br>45.0] | 30.0 [19.0,<br>45.0] |
| Primiparity |  |  |  |  |  |
| no | 158 (43.9%) | 105 (45.9%) | 137 (50.0%) | 74 (43.8%) | 474 (45.9%) |
| yes | 202 (56.1%) | 124 (54.1%) | 137 (50.0%) | 95 (56.2%) | 558 (54.1%) |
| Sex |  |  |  |  |  |
| boy | 189 (52.5%) | 125 (54.6%) | 150 (54.7%) | 98 (58.0%) | 562 (54.5%) |
| girl | 171 (47.5%) | 104 (45.4%) | 124 (45.3%) | 71 (42.0%) | 470 (45.5%) |

### Demographic factors and stool characteristics

We investigated the association between demographic factors including age, breastfeeding status, delivery mode, recent antibiotics intake and premature birth (born < 37 gestational weeks) on stool consistency and water content. As expected, individuals with parent-rated constipation exhibited lower overall water content (Figure 1B, Kruskal-Wallis test, Dunn’s test q=0.003), particularly at 6 and 14 months (Figure 1B, Kruskal-Wallis test, Dunn’s test q = 0.032, q= 0.017, respectively). Normal stool consistency was associated with 4.2 percentage point higher water content compared to constipation, adjusting for sampling age and accounting for within-subject correlation using random intercepts (p=1.2 x 10^-4^).

Our analysis revealed a declining trend in water content with increasing age (Figure 1C). On average, when accounting for the intra-individual covariance, water content was 1.6 percent lower after a year (p=4.6 x 10^-8^). Moreover, subjects who were breastfed demonstrated higher water content compared to those who were not (Figure 1D, Wilcoxon signed-rank test q = 8⋅10^-7^). However, the magnitude of the association was small, and breastfeeding vs no breastfeeding increased the water content by 1 % (linear mixed model with breastfeeding and age as fixed effects, water content as dependent variable and subject identity as random effect, p=0.047).

Overall, breastfeeding was associated with stool consistency (q = 0.00110), but delivery mode, antibiotic intake or prematurity at any specific time point or overall did not have an association. However, we observed a distinct pattern between stool consistency and age (Figure 1D). Specifically, we noted fewer occurrences of loose stool at 6 months compared to 2.5 months (Figure 1D, Dunn’s test q = 0.028). Additionally, there were more reports of gas at 2.5 months compared to 6, 14, or 30 months (Figure 1E, Dunn’s test q = 2.3 x 10-⁶, q = 8.8 x 10-^11^, q = 1.2 x 10-^7^, for 6, 14, 30 months, respectively). The prevalence of constipation was higher at 6 months compared to 2.5, 14, and 30 months (Figure 1B, Dunn’s test q = 3⋅10^-7^, q = 8⋅10^-6^, q = 0.0026 for 2.5, 14, and 30 months, respectively. Furthermore, reports on normal stool consistency were more common at 14 and 30 months compared to 2.5 months (Figure 1D, Dunn’s test q = 0.0017, q = 0.0017 for 14 and 30 months, respectively).

### Stool metabolite concentrations and stool consistency and water content

Next, we investigated whether stool metabolites were associated with stool consistency and water content. Longitudinal analyses using mixed-effects models adjusted for age, and additionally for breastfeeding status revealed that taurine and glycine-conjugated bile acids were inversely associated with constipated stool compared with normal stool (Figure 2A, Table S2). Emerging studies have shown that gut microbes can conjugate bile acids with diverse amino acids. We therefore examined whether these microbially conjugated bile acid amidates were associated with fecal water content and stool consistency. Most of these bile acids amidates-including DCA/HDCA/UDCA-Val, CA/CDCA/UDCA/HDCA-Ala, UDCA-Trp, and DCA/HDCA-Ile/Leu were negatively associated with water content (Figure 2A, Tables S3). In contrast, methionine- and alanine-conjugated bile acids were positively associated with water content (Figure 2A, Tables S3). Moreover, primary bile acids, tauro-conjugated bile acids, acetic acid, fructose, pinitol, and several other polar metabolites were positively associated with fecal water content (Figure 2B). In contrast, the branched-chain short-chain fatty acids, iso-butyric acid and iso-valeric acid were negatively associated with water content (Figure 2A, Table S3). Additionally, we also found that fecal water content was positively associated with the total sum of all SCFAs (Figure 2B; linear mixed-effects model, p = 0.1 × 10⁻¹⁷ for water content, p = 0.047 for constipation). Constipation was associated with lower levels of taurine- and glycine-conjugated bile acids, as well as the bile acid amidate CA–Asn, whereas palmitic acid (16:0) was the only metabolite significantly associated with loose stool (Figure 2C).

**Figure 2.**
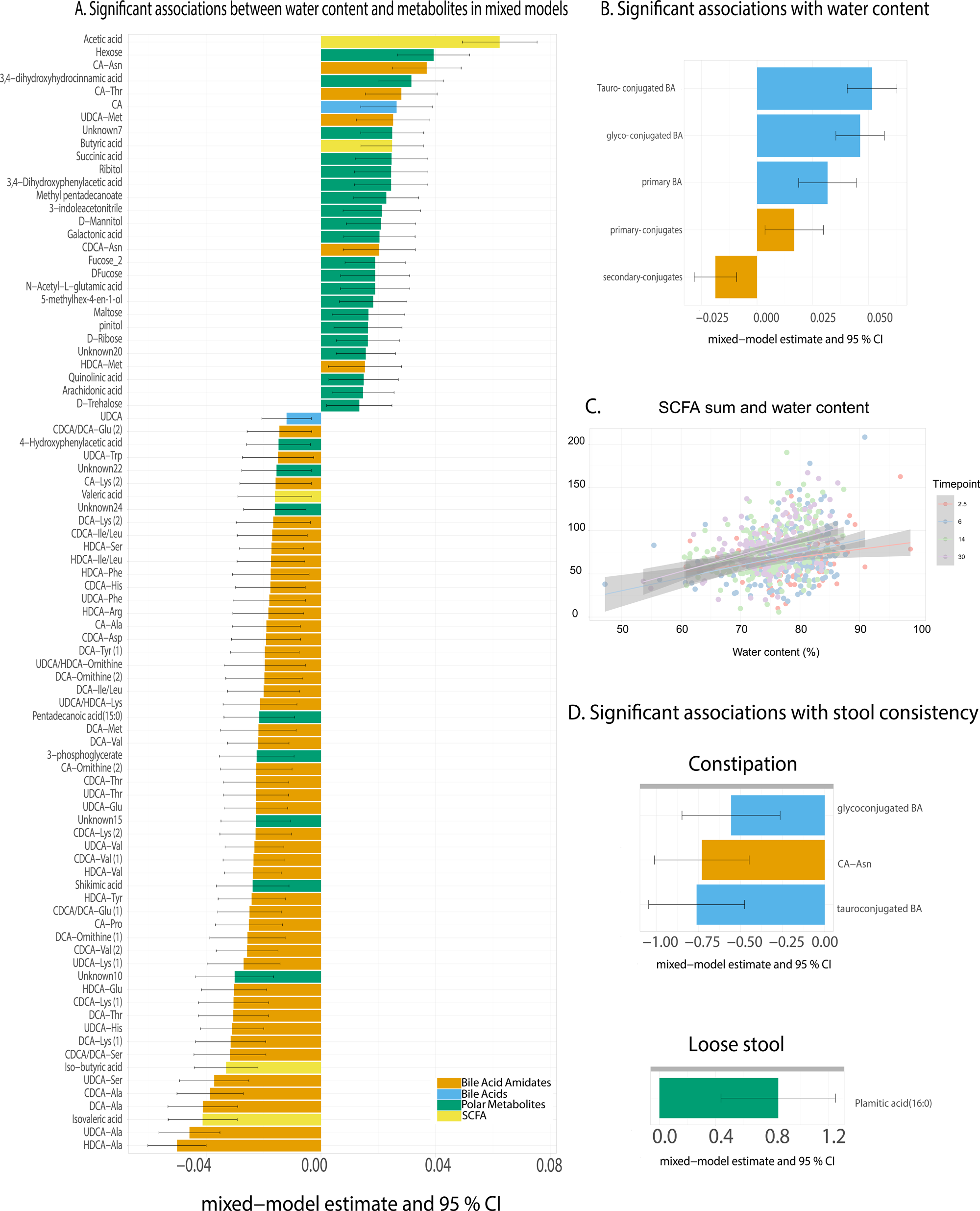
Metabolite concentrations associated with water content and constipation in mixed models with metabolite concentration as the dependent variable, stool consistency or water content as the fixed effect, any current breastfeeding as fixed effect, and child identity as random effect. A. Water content associated positive with taurine- and glycine conjugated bile acids, bile acid amidates, polar metabolites such as fucose, acetic and butyric acids. Water content associated negatively with iso-butyric and isovaleric acids, and several bile acid amidates. B. Water content associated positively with overall Bile acid and SCFA concentrations. C. The constipated individuals had lower total bile acids. The only positive association was with palmitic acid.

Given that higher stool water content and defecation frequency reflect faster intestinal transit, we next examined whether host genetic predisposition to increased stool frequency was associated with bile acid metabolites using a polygenic score (PGS). The stool-frequency PGS was positively associated with multiple bile acid amidates and negatively associated with succinic acid, independent of age but these associations did not withhold multiple testing correction (Figure S1).

### Predicted enzymatic capacity for bile acid and SCFA metabolism

Based on the results, we also examined whether predicted bacterial enzymes involved in SCFA and BA metabolism were associated with stool consistency and water content using PICRUSt2 (Phylogenetic Investigation of Communities by Reconstruction of Unobserved States). We found mainly genes involved in acetic acid metabolism, and to a lesser extent with propionic acid, butyric acid and bile acids in our data (Figure S2).

Longitudinal analyses with mixed models revealed that the abundance of SCFA-metabolizing enzymes was positively associated with constipation compared to lose stool consistency, after adjusting for age and breastfeeding. Furthermore, in a linear mixed models adjusted for age and breastfeeding as fixed effects and child identity as random effects, the abundance of acetic, butyric, and propionic acid-metabolizing bacterial enzymes was negatively associated with stool water content. Similarly, for bile acid metabolism, we found that the abundance of chologlycine hydrolase, a bile salt hydrolase primarily involved in the deconjugation of primary bile acids, was negatively associated with stool water content (Figure S3), consistent with altered bile acid deconjugation in the context of faster intestinal transit.

### Taxonomic richness, diversity and composition associate with water content and stool consistency

Next, we sought to determine what changes in microbiome were reflected with stool consistency and water content. In a mixed model, both observed richness and Shannon index associated (Figure 3A, Table S4) with water content. However, neither observed richness nor Shannon index were associated with stool consistency across time points or longitudinally (Table S4).

**Figure 3.**
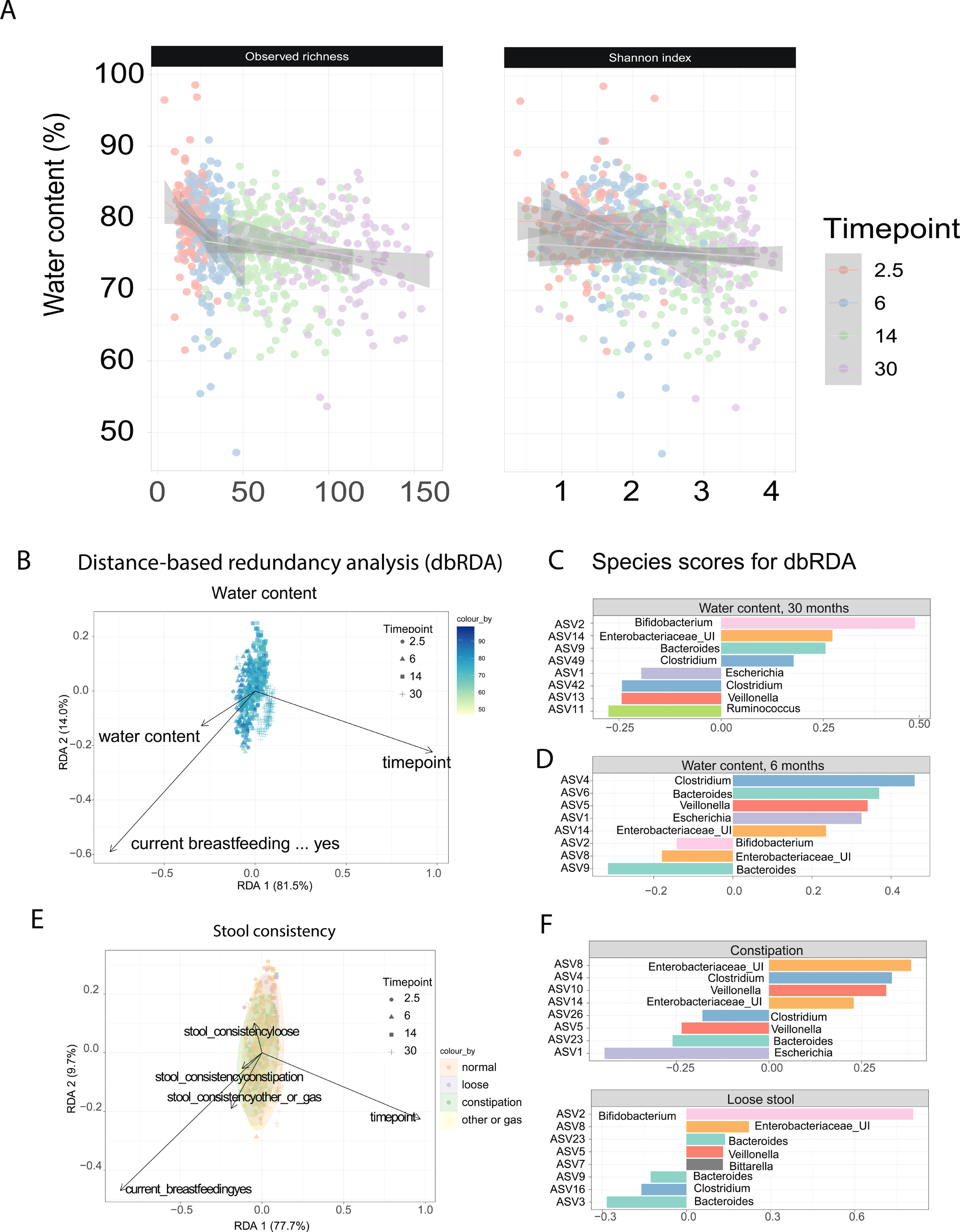
Stool consistency, water content and microbiome. A) Taxonomic richness (observed richness) and diversity (Shannon index) and water content. B. Stool consistency and community composition. db-RDA on A. water content and B. stool consistency. C. shows the ASVs that have the largest absolute coefficients in the RDA models. The bar colors represent the genus-level annotation of the ASV. Here, enterobacteriaceae UI is Enterobacteriaceae unidentified genus.

Community composition was associated with water content in distance-based redundancy analysis (Figure 3B, Table S5, Bray-Curtis dissimilarity, db-RDA PERMANOVA R^2^ = 0.006, q = 0.012) at both 6 (Table S5, R^2^ = 0.015, q = 0.012) and 30 months (Table S4, R^2^ = 0.015, q = 0.016). This association persisted after adjusting for breastfeeding in the entire dataset for 6 months. None of the subjects were breastfed for 30 months. The associations were similar with Aitchison distance, and additionally water content at 14 months was associated with water content while adjusting for breastfeeding. (Table S6, R^2^ = 0.012, q = 0.02). At 6 months, variance in water content was positively associated with amplicon sequence variants (ASVs) annotated to *Clostridium, Escherichia,* and *Veillonella*, while it was inversely associated with *Bifidobacterium* (Figure 3B). ASVs in *Bacteroides* and *Enterobacteriaceae* showed loading in both directions. Conversely, in 30-month-olds, the relative abundance of *Veillonella, Sutterella,* and *Ruminococcus* were associated with lower water content, while *Bacteroides, Bifidobacterium,* and *Enterobacteriaceae* were associated with higher water content. Community composition was associated with overall stool consistency (Figure3D-F, Table S4, PERMANOVA R^2^ = 0.003, q = 0.014), even after adjusting for breastfeeding (Table S5, PERMANOVA R^2^ = 0.041, q = 0.038). The associations remained similar while using Aitchison distance (Table S6). Stool-frequency PGS was not related to gut microbiota composition or diversity (Table S4-6).

We also performed genus-level differential abundance analysis using MaAsLin 3, adjusting for current breastfeeding status and age, with subject included as a random effect. No associations between microbial abundances and stool water content or stool consistency remained significant after multiple-testing correction (Table S7). At nominal significance (p < 0.05), stool consistency was inversely associated with Streptococcus, Alistipes, Anaerotruncus, Blautia, and Clostridium_1, and positively associated with Unidentified_Genus_6, Enterococcus, Anaeroglobus, Roseburia, and Citrobacter (Table S7). In contrast, stool-frequency polygenic score (PGS) was significantly associated with Acidaminococcus abundance and prevalence (Figure S7).

## Discussion

We show in infants and small children that both stool microbiota and metabolome link with the stool consistency and the total water content. This corroborates earlier findings indicating that stool consistency and gut transit time correlate with microbiota and metabolism in adults [2, 14, 43]. However, data especially from infants and with longitudinal study design are scarce. Our findings indicate more extensive microbial metabolism in low-moisture stool and in constipated children. Moreover, our findings also underscore the potential of using the total water content as an objective measure in microbiota studies, particularly during early infancy when stool consistency vary considerably. We also found agreement between the total stool water content and parental crude reports of constipation.

Regarding the metabolic features, we show that the associations between metabolites and low stool water content and parent-report of constipation are qualitatively similar: children with constipation and lower water content also showed decreased concentrations of host-conjugated bile acids (glycine and taurine-bound). Gut microbes regulate human bile acid metabolism [44]. Primary bile acids are synthesized and conjugated in the liver, stored in the gallbladder, secreted in the intestine after meal consumption, and later modified (de-conjugated or again re-conjugated) by microbes in the gut before reabsorption (95%) or excretion (5%) [45]. Thus, the decreased level of host-conjugated BAs in egested constipated stool suggests that longer transit time allows for more extensive bacterial metabolism of bile acids in the gut[43]. Moreover, bacteria can then conjugate bile acids with other amino acids and produce novel conjugated bile acid amidates. In line, many microbially conjugated bile acid amidates were negatively associated with water content as well as the polygenic score for defecation frequency. Aberrant bowel movement frequencies have been linked to constipation and alterations in microbially derived metabolites in the circulation [46]. This again indicates that longer transit time allows for more microbially driven re-conjugation of bile acids. This might have health implications, since biliary excretion of xenobiotics is an important elimination route, and deconjugation might delay clearance of substances such as estrogens [47].

Besides bile acids, constipation and lower water content were associated with less SCFA, specifically acetic acid, while branched short-chain fatty acids (BCFAs) concentrations were inversely associated with stool water content at 14 and 30 months. In fact, SCFAs, primarily acetate, propionate, and butyrate, are end products of fermentation of complex carbohydrates such as dietary fiber [48, 49], while increased BCFAs reflect ongoing putrefaction or/and proteolytic activities in the colon. We hypothesize that prolonged colonic transit time depleted available carbohydrates for fermentation which consequently induced proteolytic activity, regarded as an alternative pathway for carbon and nitrogen, and an energy source for the microbes in the colon. Interestingly, another study showed that by reducing gut transit time with drugs, the concentrations of total SCFAs could be increased [50]. Our finding is consistent with similar works in adults, where lower water content or longer gut transit time showed to shift in colonic metabolism towards proteolytic activity [14, 43, 50–52]. Roager *et al*. found that longer transit time was associated with higher levels of protein-derived molecules in the urine such as p-cresol-sulfate and phenylacetylglutamine, indicating a shift from saccharolytic to proteolytic activity in the colon [14]. Also, Zhang *et al*. found that the Bristol stool form scale scores were found to negatively associate with the levels of fecal amino acids, suggesting increased protein catabolism in irritable bowel syndrome patients [53]. In adults, metabolites from protein catabolism have been implicated in atherosclerotic cardiovascular diseases and colon cancer [54–56]. Hence our observation that higher BCFA concentrations were associated with constipation might have further health-implications, although this has not yet been studied in infants or small children.

The predicted metabolic capacity for SCFA and BA showed that genes related to SCFA, mainly acetic acid, metabolism were negatively associated with water content and increased in constipated individuals. Although 16s rRNA gene sequencing has limited resolution in predicting functional capacity, and bile acid metabolism had presently limited coverage in the database, we could show that the microbiome relates with proxies of transit time not only on compositional level, but also on predicted functional properties. Interestingly, bile salt hydrolase was negatively associated with water content. Bile salt hydrolase possesses dual functions in bile acid metabolism; it has been shown to both deconjugate as well re-conjugate these novel microbial bile acids amidate [57]. Hence it is plausible that faster transit time relates to lower abundance of bile salt hydrolase.

Fluctuations in stool consistency correlate with microbiota taxonomic richness, and community composition[2, 17, 43, 58]. Vandeputte et al. (2016) found that firmer stools in Bristol stool form scale were associated with a higher Bacteroidetes:Firmicutes ratio and elevated levels of *Akkermansia* and *Methanobrevibacter* whereas loose stools were associated with lower taxonomic richness in adults [2]. In line, our study in infants and small children reveals that stool consistency associates with the bacterial community composition, particularly through variations in *Bacteroides, Bifidobacterium, Ruminococcus,* and *Clostridium* abundances. We also found an ASV in *Bacteroides* genus associated with the water content-related community composition positively at 30 months and negatively at 6 months, potentially due to changing diet and overall gut microbiome composition.

Similarly, a previous study reported a different direction in the association between stool consistency and microbiota composition during infancy and childhood [21]. Intriguingly, Kwiatkowska *et al.* 2024 reported that functional constipation in three-year-olds was associated with depletion of rare genera, such as butyrate-producer *Subdoligranulum [59].* We observed that constipated individuals had more homogenous community composition compared to other stool consistency groups, and the lack of rare taxa could partly explain our observation, however, we did not directly assess prevalence or abundance of rare taxa. Overall, these findings align with previous research on the relationship between stool consistency and microbiota, although differences in associated taxonomic groups may result from gut microbiota alterations across different life stages [60].

We expect that our findings may reflect the influence of longer colonic transit time on intestinal microbial communities potentially via selectively favoring certain microbes and their activity and levels of microbe-regulated metabolites[58]. However, other mechanisms may underline the links between microbiota and gut motility. Although we observed concurrent associations and we did not study predictive capacity of gut microbiome on later markers of transit time, it should be noted that early life microbiota may affect the development of enteric nervous system and consequently (later) gut motility and secretions [61–64]. Although we studied generally healthy sample, our results highlight a need to study whether similar microbiota and microbially regulated metabolite signatures could be seen in pathological context, such as in disorders of gut-brain interaction in older children [59, 65–67].

Our findings of parent-reported stool consistency, water content, child age and breastfeeding are in line with earlier literature and clinics. First, the information that parent-reported constipation associated with objective measure of lower water content may be clinically relevant to support future research to avoid invasive examination techniques of colonic transit time [68, 69]. Second, the association between breastfeeding and stool consistency and water content is in line with general clinical observation that breastfed infants seldom suffer from constipation [70], and prolonged breastfeeding may protect from functional constipation[71]. Moreover, parental reports of gas were prevalent at 2.5-months-of-age when over half of the infants were breastfed and they typically swallow air during breastfeeding which leads to gas problems [70] . Third, constipation becomes more prevalent at 6 months that aligns well with the introduction of solid foods and reduction of breastfeeding. Our observation that water content gradually decreases by age likely reflects the normative prolonging of gut transit time by age [72, 73]

Our study has limitations. We used crude parental report of child stool consistency, which is not as refined as the Bristol stool scale, and this may have implications on comparability or generalizability of our results but still, it was associated with the objective measure of total stool water content. Stool water content is different from *e.g.,* gastrointestinal transit time assessment with radio-opaque markers, but we studied vulnerable study subjects of healthy infants and young children, where utilizing invasive measuring techniques has ethical constrains. Utilizing stool samples does not allow proper inference of absorption of the microbiota-produced metabolites in the whole gut. Serum or urinary metabolome analysis may have given greater insight into the systemic alterations related to the saccharolytic *vs.* proteolytic fermentation and enterohepatic circulation. Moreover, one can think that extremely low water content in the faeces may alter the DNA extraction success, which may subsequently lead to changes in the microbiota composition. However, we are not aware of methodological studies addressing this.

## Conclusion

In summary, our findings in infants and small children support the concept that stool consistency and total water content are associated with gut microbiota composition and metabolic activity. Our result suggests metabolic shifts from saccharolytic to proteolytic activity and decreased concentrations of host-conjugated bile acids and increased concentrations of microbially conjugated bile acids related to constipation and lower water content, providing insights into gut health and function. Stool consistency and water content were further linked with predicted SCFA and bile acid metabolizing capacity by the gut microbes. We found microbiota community composition to be associated with stool consistency, and these varied partly by age.

## Author Contributions

Conception or design of the work: AKA, SL, AMD, TB. Acquisition, data analysis, or interpretation of data: AP, TP, KK, PCD, LK, HK, HI, MK, LL, ML, MO, AMD, AKA, SL, EM. SL and AKA wrote the manuscript. All authors critically reviewed and approved the final manuscript. All authors have approved the submitted revision.

## Acknowledgments

Finnbrain Birth cohort Study (HK) has been funded by Research Council of Finland (grant numbers 253270, 134950), Jane and Aatos Erkko Foundation, as well as Signe and Ane Gyllenberg Foundation. LK was funded by the Research Council of Finland (grant number 308176 and 325292), Yrjö Jahnsson Foundation (6847, 6976), Signe and Ane Gyllenberg Foundation, Finnish State Grants for Clinical Research (P3654), Jalmari and Rauha Ahokas Foundation, and Waterloo Foundation (2110-3601). AKA was supported by Yrjö Jahnsson Foundation, Psychiatry Research Foundation, Emil Aaltonen Foundation, Brain Foundation, Instrumentarium Science Foundation, Signe and Ane Gyllenberg Foundation, Duodecim Finnish Medical Society, Juho Vainio Foundation and the Research Council of Finland (grant number 347640). S.L was supported by the Research Council of Finland (decision number 363417). LL was supported by Research Council of Finland (grant number 330887). AMD has been funded by the Waterloo foundation and Research Council of Finland (347924). Further support was received by the “Inflammation in human early life: targeting impacts on life-course health” (INITIALISE) consortium funded by the Horizon Europe Program of the European Union under Grant Agreement 101094099 (to MO), and InFLAMES Flagship Programme of the Research Council of Finland (decision number: 337530). AKA and HI were supported by the Signe and Ane Gyllenberg Foundation (grant no. 6273). HI received funding from the Finnish Cultural Foundation (grant no. 00230482), the Finnish Foundation for Cardiovascular Research, the Diabetes Research Foundation, and the Doctoral Program in Clinical Research at the University of Turku. We thank the FinnBrain Birth Cohort Study participants and personnel, Turku Metabolomics Center for the assistance and resources in the analyses of metabolites. We also thank Paulo Wender P Gomes for his excellent suggestions with the graphics. We like to acknowledge the Maternal and Pediatric Precision In Therapeutics (MPRINT) program, funded by NICHD P50 award (P50HD106463). We would like to acknowledge that the current version of this manuscript is available as a preprint on bioRxiv [74].

## Appendix A. Supplementary Information

### Supplementary figures

S1 PGS associations with metabolites

S2 Metabolizing capacity of predicted ECs found in the data set

S3 EC abundances associations with water content and stool consistency

S4 PGS association with Acidaminococcus

### Supplementary Tables

S1 Sample sizes for available data types for both stool consistency and water content analyses. Sample sizes listed with only the main outcome variable, i.e. all, as well as with available covariate data (time point and current breastfeeding), i.e. covariates.

S2 Metabolite associations with stool consistency in a mixed-model adjusted for breastfeeding

S3 Metabolite associations with water content in a mixed-model adjusted for breastfeeding

S4 Mixed model estimate, p-, and q-values for taxonomic diversity, water content, stool consistency and PGS as predictors

S5 PERMANOVA using Bray-Curtis dissimilarity results for water content, stool consistency and PGS

S6 PERMANOVA using Aitchison distance results for water content, stool consistency and PGS

S7 All MaAsLin3 results for the main independent variables

## Data and code availability

Due to Finnish national legislation and study participant consents, the individual-level metadata cannot be made available online, but data can potentially be shared with Research Agreement as part of research collaboration. Requests for collaboration can be sent to the Board of the FinnBrain Birth Cohort Study; please contact Linnea Karlsson. The R scripts for data analyses can be found in Zenodo (DOI: 10.5281/zenodo.11222840).

## References

1. Lyon, L., ’All disease begins in the gut’: was Hippocrates right? Brain, 2018. 141(3): p. e20.

2. Vandeputte, D., et al., Stool consistency is strongly associated with gut microbiota richness and composition, enterotypes and bacterial growth rates. Gut, 2016. 65(1): p. 57–62.

3. Tigchelaar, E.F., et al., Gut microbiota composition associated with stool consistency. Gut, 2016. 65(3): p. 540–2.

4. Hadizadeh, F., et al., Stool frequency is associated with gut microbiota composition. Gut, 2017. 66(3): p. 559–560.

5. Deutsch, L. and B. Stres, The Importance of Objective Stool Classification in Fecal 1H-NMR Metabolomics: Exponential Increase in Stool Crosslinking Is Mirrored in Systemic Inflammation and Associated to Fecal Acetate and Methionine. Metabolites, 2021. 11(3).

6. Yap, C.X., et al., Autism-related dietary preferences mediate autism-gut microbiome associations. Cell, 2021. 184(24): p. 5916–5931.e17.

7. Fan, Y. and O. Pedersen, Gut microbiota in human metabolic health and disease. Nat Rev Microbiol, 2021. 19(1): p. 55–71.

8. Lamichhane, S., et al., Dysregulation of secondary bile acid metabolism precedes islet autoimmunity and type 1 diabetes. Cell Rep Med, 2022. 3(10): p. 100762.

9. Fromentin, S., et al., Microbiome and metabolome features of the cardiometabolic disease spectrum. Nat Med, 2022. 28(2): p. 303–314.

10. Talmor-Barkan, Y., et al., Metabolomic and microbiome profiling reveals personalized risk factors for coronary artery disease. Nat Med, 2022. 28(2): p. 295–302.

11. Lamichhane, S., et al., Gut metabolome meets microbiome: A methodological perspective to understand the relationship between host and microbe. Methods, 2018. 149: p. 3–12.

12. Zierer, J., et al., The fecal metabolome as a functional readout of the gut microbiome. Nat Genet, 2018. 50(6): p. 790–795.

13. Knight, R., et al., Best practices for analysing microbiomes. Nat Rev Microbiol, 2018. 16(7): p. 410–422.

14. Roager, H.M., et al., Colonic transit time is related to bacterial metabolism and mucosal turnover in the gut. Nat Microbiol, 2016. 1(9): p. 16093.

15. Hasan, N. and H. Yang, Factors affecting the composition of the gut microbiota, and its modulation. PeerJ, 2019. 7: p. e7502.

16. Asnicar, F., et al., Blue poo: impact of gut transit time on the gut microbiome using a novel marker. Gut, 2021. 70(9): p. 1665–1674.

17. Procházková, N., et al., Advancing human gut microbiota research by considering gut transit time. Gut, 2023. 72(1): p. 180–191.

18. Falony, G., et al., Population-level analysis of gut microbiome variation. Science, 2016. 352(6285): p. 560-4.

19. Schindlbeck, N.E., A.G. Klauser, and S.A. Müller-Lissner, [Measurement of colon transit time]. Z Gastroenterol, 1990. 28(8): p. 399–404.

20. Degen, L.P. and S.F. Phillips, How well does stool form reflect colonic transit? Gut, 1996. 39(1): p. 109–13.

21. Jokela, R., et al., Sources of gut microbiota variation in a large longitudinal Finnish infant cohort. EBioMedicine, 2023. 94: p. 104695.

22. Karlsson, L., et al., Cohort Profile: The FinnBrain Birth Cohort Study (FinnBrain). Int J Epidemiol, 2018. 47(1): p. 15–16j.

23. Aatsinki, A.-K., et al., Dynamics of Gut Metabolome and Microbiome Maturation during Early Life. medRxiv, 2023: p. 2023.05.29.23290441.

24. Rintala, A., et al., Early fecal microbiota composition in children who later develop celiac disease and associated autoimmunity. Scand J Gastroenterol, 2018. 53(4): p. 403–409.

25. Callahan, B.J., et al., DADA2: High-resolution sample inference from Illumina amplicon data. Nat Methods, 2016. 13(7): p. 581–3.

26. Quast, C., et al., The SILVA ribosomal RNA gene database project: improved data processing and web-based tools. Nucleic Acids Res, 2013. 41(Database issue): p. D590-6.

27. Thapa, K., et al., Melanocortin 1 receptor regulates cholesterol and bile acid metabolism in the liver. Elife, 2023. 12.

28. Kråkström, M., et al., Dynamics of the Lipidome in a Colon Simulator. Metabolites, 2023. 13(3): p. 355.

29. Huovinen, V., et al., Infant gut microbiota and negative and fear reactivity. Dev Psychopathol, 2023: p. 1–16.

30. Wiese, M., et al., CoMiniGut-a small volume in vitro colon model for the screening of gut microbial fermentation processes. PeerJ, 2018. 6: p. e4268.

31. Lamichhane, S., et al., Circulating metabolites in progression to islet autoimmunity and type 1 diabetes. Diabetologia, 2019. 62(12): p. 2287–2297.

32. Douglas, G.M., et al., PICRUSt2 for prediction of metagenome functions. Nat Biotechnol, 2020. 38(6): p. 685–688.

33. Kanehisa, M., et al., KEGG: new perspectives on genomes, pathways, diseases and drugs. Nucleic Acids Res, 2017. 45(D1): p. D353–d361.

34. Shimizu, N., et al., A novel RNA splicing mutation in Japanese patients with Wilson disease. Biochem Biophys Res Commun, 1995. 217(1): p. 16–20.

35. Loh, P.R., et al., Reference-based phasing using the Haplotype Reference Consortium panel. Nat Genet, 2016. 48(11): p. 1443–1448.

36. Browning, B.L. and S.R. Browning, Genotype Imputation with Millions of Reference Samples. Am J Hum Genet, 2016. 98(1): p. 116–26.

37. Bonfiglio, F., et al., GWAS of stool frequency provides insights into gastrointestinal motility and irritable bowel syndrome. Cell Genom, 2021. 1(3): p. None.

38. Ge, T., et al., Polygenic prediction via Bayesian regression and continuous shrinkage priors. Nat Commun, 2019. 10(1): p. 1776.

39. Ernst F, S.S., Borman T, Lahti L. mia: Microbiome analysis. R package version 1.12.0, https://github.com/microbiome/mia. 2024.

40. McCarthy, D.J., et al., Scater: pre-processing, quality control, normalization and visualization of single-cell RNA-seq data in R. Bioinformatics, 2017. 33(8): p. 1179–1186.

41. Dixon, P., VEGAN, a package of R functions for community ecology. Journal of Vegetation Science, 2003. 14(6): p. 927–930.

42. Pinheiro J, B.D., R Core Team (2023). nlme: Linear and Nonlinear Mixed Effects Models. R package version 3.1–164, https://CRAN.R-project.org/package=nlme.

43. Müller, M., et al., Distal colonic transit is linked to gut microbiota diversity and microbial fermentation in humans with slow colonic transit. Am J Physiol Gastrointest Liver Physiol, 2020. 318(2): p. G361–g369.

44. Mohanty, I., et al., The changing metabolic landscape of bile acids - keys to metabolism and immune regulation. Nat Rev Gastroenterol Hepatol, 2024.

45. Guzior, D.V. and R.A. Quinn, Review: microbial transformations of human bile acids. Microbiome, 2021. 9(1): p. 140.

46. Johnson-Martínez, J.P., et al., Aberrant bowel movement frequencies coincide with increased microbe-derived blood metabolites associated with reduced organ function. Cell Rep Med, 2024. 5(7): p. 101646.

47. Hu, S., et al., Gut microbial beta-glucuronidase: a vital regulator in female estrogen metabolism. Gut Microbes, 2023. 15(1): p. 2236749.

48. Lamichhane, S., et al., Metabolic Fate of (13)C-Labeled Polydextrose and Impact on the Gut Microbiome: A Triple-Phase Study in a Colon Simulator. J Proteome Res, 2018. 17(3): p. 1041–1053.

49. Kråkström, M., et al., Dynamics of the Lipidome in a Colon Simulator. Metabolites, 2023. 13(3).

50. El Oufir, L., et al., Relations between transit time, fermentation products, and hydrogen consuming flora in healthy humans. Gut, 1996. 38(6): p. 870–7.

51. Lewis, S.J. and K.W. Heaton, Increasing butyrate concentration in the distal colon by accelerating intestinal transit. Gut, 1997. 41(2): p. 245–51.

52. Gabriele, S., et al., Slow intestinal transit contributes to elevate urinary p-cresol level in Italian autistic children. Autism Res, 2016. 9(7): p. 752–9.

53. Zhang, W.X., et al., Altered profiles of fecal metabolites correlate with visceral hypersensitivity and may contribute to symptom severity of diarrhea-predominant irritable bowel syndrome. World J Gastroenterol, 2019. 25(43): p. 6416–6429.

54. Wang, Z., et al., Gut flora metabolism of phosphatidylcholine promotes cardiovascular disease. Nature, 2011. 472(7341): p. 57-63.

55. Nemet, I., et al., A Cardiovascular Disease-Linked Gut Microbial Metabolite Acts via Adrenergic Receptors. Cell, 2020. 180(5): p. 862–877.e22.

56. Al Hinai, E.A., et al., Modelling the role of microbial p-cresol in colorectal genotoxicity. Gut Microbes, 2019. 10(3): p. 398–411.

57. Rimal, B., et al., Bile salt hydrolase catalyses formation of amine-conjugated bile acids. Nature, 2024. 626(8000): p. 859-863.

58. Minnebo, Y., et al., Gut microbiota response to in vitro transit time variation is mediated by microbial growth rates, nutrient use efficiency and adaptation to in vivo transit time. Microbiome, 2023. 11(1): p. 240.

59. Kwiatkowska, M., et al., The oral cavity and intestinal microbiome in children with functional constipation. Sci Rep, 2024. 14(1): p. 8283.

60. Martino, C., et al., Microbiota succession throughout life from the cradle to the grave. Nat Rev Microbiol, 2022. 20(12): p. 707–720.

61. Hung, L.Y., et al., Antibiotic exposure postweaning disrupts the neurochemistry and function of enteric neurons mediating colonic motor activity. Am J Physiol Gastrointest Liver Physiol, 2020. 318(6): p. G1042–g1053.

62. Griffiths, J.A., et al., Peripheral neuronal activation shapes the microbiome and alters gut physiology. Cell Rep, 2024. 43(4): p. 113953.

63. Caputi, V., et al., Antibiotic-induced dysbiosis of the microbiota impairs gut neuromuscular function in juvenile mice. Br J Pharmacol, 2017. 174(20): p. 3623–3639.

64. De Vadder, F., et al., Gut microbiota regulates maturation of the adult enteric nervous system via enteric serotonin networks. Proc Natl Acad Sci U S A, 2018. 115(25): p. 6458–6463.

65. de Moraes, J.G., et al., Fecal Microbiota and Diet of Children with Chronic Constipation. Int J Pediatr, 2016. 2016: p. 6787269.

66. Hofman, D., et al., Faecal Microbiota in Infants and Young Children with Functional Gastrointestinal Disorders: A Systematic Review. Nutrients, 2022. 14(5).

67. Yuan, Y., et al., Characteristics of the Cajal interstitial cells and intestinal microbiota in children with refractory constipation. Microb Pathog, 2023. 184: p. 106373.

68. Sharif, H., et al., Imaging Measurement of Whole Gut Transit Time in Paediatric and Adult Functional Gastrointestinal Disorders: A Systematic Review and Narrative Synthesis. Diagnostics (Basel), 2019. 9(4).

69. Popescu, M. and M. Mutalib, Bowel transit studies in children: evidence base, role and practicalities. Frontline Gastroenterol, 2022. 13(2): p. 152–159.

70. Salvatore, S., et al., Review shows that parental reassurance and nutritional advice help to optimise the management of functional gastrointestinal disorders in infants. Acta Paediatr, 2018. 107(9): p. 1512–20.

71. Motoki, N., et al., Impact of breastfeeding during infancy on functional constipation at 3 years of age: the Japan Environment and Children’s Study. Int Breastfeed J, 2023. 18(1): p. 57.

72. Bowles, A., et al., Specific aspects of gastro-intestinal transit in children for drug delivery design. Int J Pharm, 2010. 395(1-2): p. 37–43.

73. Tran, D.L. and P. Sintusek, Functional constipation in children: What physicians should know. World J Gastroenterol, 2023. 29(8): p. 1261–1288.

74. Aatsinki, A.-K., et al., Fecal microbiota and metabolite composition associates with stool consistency in young children. bioRxiv, 2024: p. 2024.06.05.597641.

