## Supplementary Figures for "Early life gut microbial and metabolic shifts reflected in stool consistency- a gut transit time proxy"

S1 PGS associations with metabolites

S2 Metabolizing capacity of predicted ECs found in the data set

S3 EC abundances associations with water content and stool consistency

S4 PGS association with Acidaminococcus

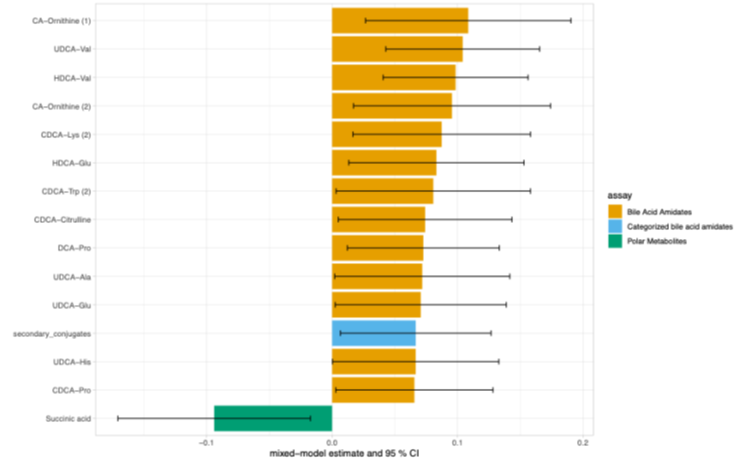

Figure S1. Polygenic score for defecation frequency associated with several secondary bile acid amidates as well as succinic acid with nominal p<0.05.

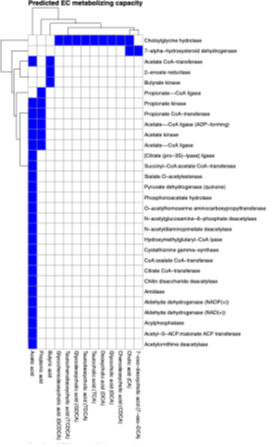

Figure S2. The heatmap indicates in blue the SCFA or BA that the given EC metabolizes. Most of the predicted anzymes were metabolizing acetic acid, but few ECs also metabolize other SCFA and BA.

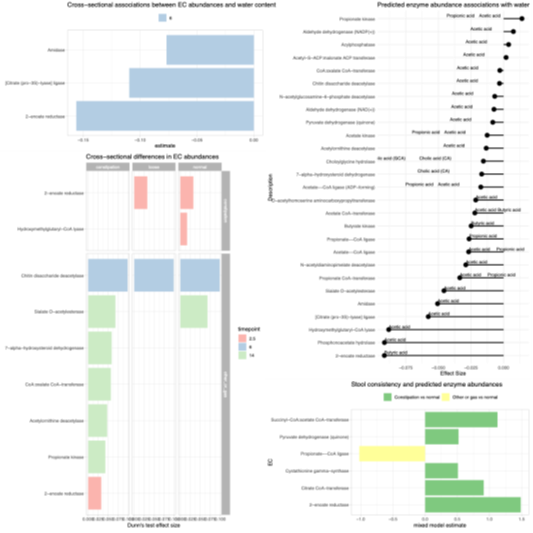

Figure S3. Microbial predicted enzymatic capacity for SCFA or BA metabolism associated with water content and stool consistency.

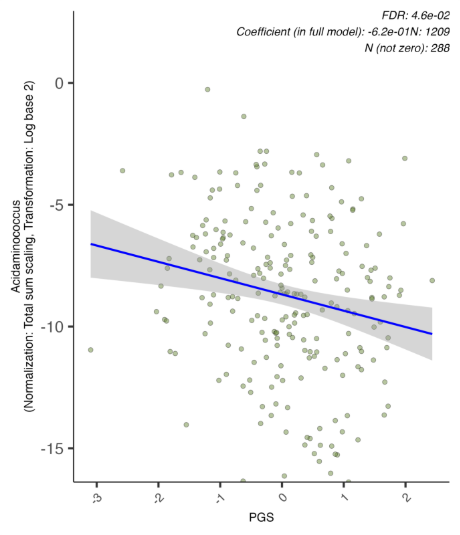

Figure S4. Acidaminococcus abundance and prevalence were associated with the PGS in MaAsLin 3.
